# Evidence that SPIO Chain Formation is Essential for High-Resolution MPI

**DOI:** 10.1101/2022.11.27.518061

**Authors:** Caylin Colson, KL Barry Fung, Jacob Bryan, Zhi Wei Tay, Benjamin D. Fellows, Chinmoy Saayuja, Renesmee Kuo, Prashant Chandrasekharan, Steven M. Conolly

## Abstract

Magnetic Particle Imaging (MPI) is a noninvasive imaging modality that exploits the saturation properties of superparamagnetic iron oxide particles (SPIOs). A major thrust of MPI research aims to sharpen the magnetic resolution of biocompatible SPIOs, which will be crucial for affordable and safe clinical translation. We recently reported on a new class of MPI tracers —called superferromagnetic iron oxide nanoparticles (SFMIOs) — which offer much sharper magnetic saturation curves. SFMIOs experimentally demonstrate 10-fold improvement in *both* resolution and sensitivity. However, superferromagnetism is a relatively unexplored branch of physics and the nanoscale physics and dynamics of SFMIOs remain a mystery. Here we show experimentally that chaining of SPIOs can explain SFMIO’s boost in SNR and resolution. We show how concentration, viscosity, transmit amplitude, and pre-polarization time can all affect SPIO chain formation and SFMIO behavior. These experiments will inform strategies on SFMIO chemical synthesis as well as SFMIO data acquisition pulse sequences.

## I. Introduction

**M**AGNETIC PARTICLE IMAGING (MPI) is a radiation-free, non-invasive imaging modality that was developed by Gleich and Weizenecker [1]. MPI is a completely different modality from Magnetic Resonance Imaging (MRI) and it requires a special-purpose hardware scanner. MPI images the saturation properties of superparamagnetic iron oxide nanoparticles (SPIOs). MRI images the precession of the water proton’s nuclear moments [2]. SPIOs are sometimes employed as a T1 or T2* contrast agent in MRI. In MPI, we detect the nonlinear signature (e.g., harmonics) from a flip of SPIOs. Importantly, there are zero harmonics from surrounding human or mammalian tissue, meaning MPI has superb contrast-to-noise ratio (CNR). The MPI signal is perfectly linear in the amount of iron within a voxel [3]–[5]. The electronic magnetization of SPIOs is 22 million times greater than the nuclear magnetization of ^1^H at 7T; so MPI scanners can measure even 5 nanograms of iron per voxel (roughly 200 cells) [2], [3]. Finally, because typical MPI scans use drive fields with frequencies in the very low frequency (VLF) regime (∼20 kHz) [6], there is no signal attenuation with depth inside the animal or human. Depth attenuation is a challenge for many imaging methods, including optical, ultrasound, and nuclear medicine. MPI’s unique physics makes it an ideal tracer imaging method with applications in angiography, cell tracking, and perfusion imaging [7]–[17].

Philips [18] and a few labs have attempted to scale up MPI from pre-clinical to the much bigger bore size for human scanners necessary for clinical imaging [19], [20]. The main challenges include cost and human safety constraints (dB/dt and SAR, [2], [6] of whole-body gradient fields and drive fields necessary for high-resolution MPI [21], [22]. Gradient costs scale quadratically with gradient strength. A pre-clinical MPI murine scanner using ferucarbotran nanoparticles has a spatial resolution of 1.5 mm using a 7 T/m gradient with a 4 cm bore radius [3], [11]. Hence, a human-sized MPI scanner operating with a 1.0 T/m gradient using ferucarbotran would offer roughly 11.0 mm spatial resolution, which is not competitive with MRI (1 mm), CT (1 mm) or Nuclear Medicine (∼3 – 5 mm). While deconvolution approaches could boost this resolution somewhat, it is clear we need a higher resolution MPI tracer for human applications.

The resolution in an MPI image is a complex research topic to be sure, but the undeconvolved resolution using either System Matrix or X-space reconstruction is simply the ratio of the magnetic resolution of the nanoparticle to the gradient strength [21], [22]. All MPI scans today use superparamagnetic iron oxide nanoparticles (SPIOs) as the tracer. An ensemble of SPIOs show no remanence, meaning there is no net magnetization observed when the applied magnetic field is zero. Because the SPIOs are too far apart to interact at the concentrations employed in MPI, they respond only to the applied magnetic fields. This is the “non-interacting” assumption. Here we explore the advantages and tradeoffs of interacting SPIOs, which we are calling superferromagnetic tracers.

The steepness of a nanoparticle’s M-H curve is responsible for the magnetic resolution measured. If the applied magnetic field flips from positive to negative magnetization, so does the SPIO’s magnetization. This flip creates a change in magnetization which is inductively sensed, allowing the change in magnetization to create a pulse in the received signal. The speed of said flip and the width of the pulse in the received signal depend on physical properties of the nanoparticles. Previous work to optimize SPIO size and magnetic properties have only focused on individual nanoparticles, not assemblies [26], [28].

Recent work by Tay, et al. [29] has shown that assemblies of chain-like SPIOs, or superferromagnetic iron oxide nanoparticles (SFMIOs) can cause order-of-magnitude enhancement of both resolution and sensitivity, as seen in Fig. 1. SFMIOs are iron oxide nanoparticles that would be superparamagnetic if they weren’t interacting with one another [30]–[32]. Superferromagnetic nanoparticles flip from negative to positive saturation much faster than superparamagnetic nanoparticles. In addition to the applied magnetic field, the superferromagnetic nanoparticles are affected by one another’s induced magnetizations, enabling a faster flipping.

**Fig. 1.**
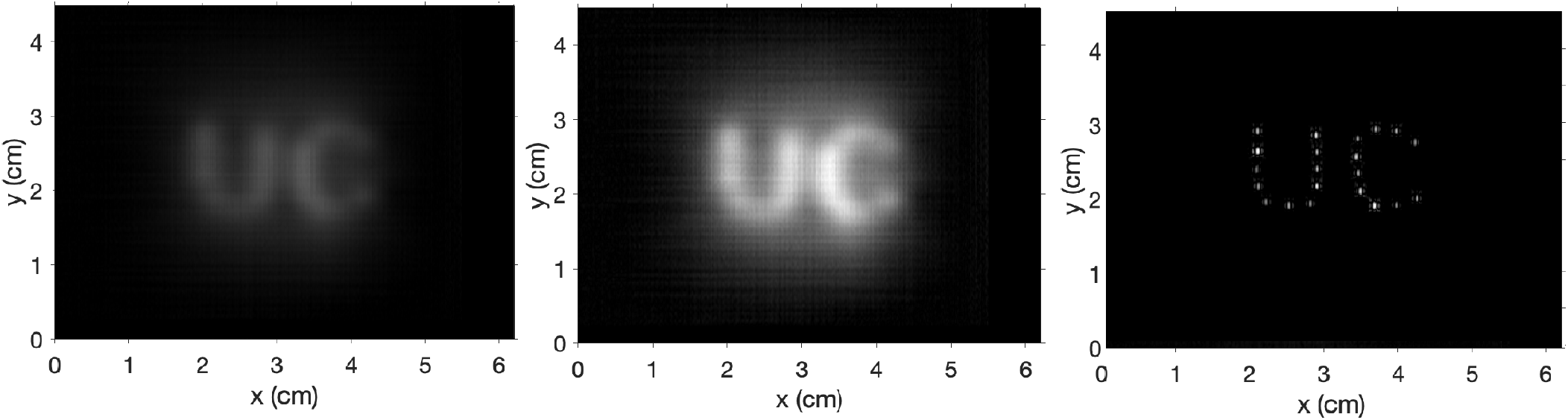
MPI Images of a UC phantom. The images in the left and middle shows SPIOs (Vivotrax), and the right image shows SFMIO tracers synthesized by our lab. The oval shape of the PSF is as predicted in [24], [25]; The normal component of the 3D MPI PSF has a spatial resolution 2.3x greater than the tangential component. The left and right images were SNR normalized; the left was then increased so that it would be more visible. The middle image is peak normalized. The measured spatial resolution (FWHM) in a 6.3 T/m gradient was ***≈***1.5 mm for SPIOs vs 300 ***µ***m for SFMIOs. This demonstrates SFMIOs can improve resolution by 5-fold and offer a 17-fold boost in SNR.

In this work, we show experimental data that supports the hypothesis that inter-particle interactions and formation of chains is responsible for the marked improvement in MPI resolution and sensitivity seen with SFMIOs. We find that SFMIO behavior depends on the following conditions: local concentration, solvent viscosity, transmit amplitude, and the length of time they are exposed to a magnetic field. Understanding the physics of SFMIOs is essential for biocompatible encapsulation and pulse sequence design.

## II. Theory

### A. X-space Magnetic Particle Imaging with Superparamagnetic Iron Oxide Nanoparticles

MPI measures the magnetization of superparamagnetic iron oxide nanoparticles in response to an applied magnetic field, which, in a 1D X-space model, is proportional to the Langevin function. Because these particles are superparamagnetic and their magnetic moments instantaneously align with an applied field, a changing applied magnetic field creates changing magnetization of the nanoparticles. The changing magnetization is measured inductively, so MPI’s 1D point spread function is proportional to the derivative of the Langevin Function.

The spatial resolution (FWHM) of the MPI equation depends only on the strength of *G*, the Gradient field [*T/m*], and the magnetic properties of the nanoparticles themselves, *k* [*m/A*], where, for a single domain magnetic nanoparticle, 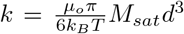. The magnetic resolution of an MPI scan using traditional SPIO particles is inversely proportional to the strength of the gradient field and inversely proportional to the cube of the magnetic core diameter. Intuitively, one would keep increasing the magnetic core diameter of SPIOs to optimize MPI resolution [21], [22]. However, Tay et al. [23] found that, after the nanoparticles reached a certain magnetic core diameter, the magnetic relaxation of the particles started to dominate, leading to blurring in the MPI image. The larger the magnetic core of a superparamagnetic particle, the further away it is from the single-domain particle description that describes superparamagnetism.

The traditional Langevin physics used to describe superparamagnetic nanoparticles has a few assumptions that must be met to accurately predict behavior. 1) A superparamagnetic particle must be small enough to be a single magnetic domain. 2) When the applied magnetic field is zero, the net magnetization of the nanoparticles should also be zero. 3) The magnetization of an SPIO is described by the Langevin Function. The advantage in using superparamagnetic nanoparticles for MPI is both in their ‘instantaneous’ flip from negative to positive saturation and in their nonlinear magnetic saturation.

### B. Interacting Superparamagnetic Nanoparticles

Recent work by Tay, et al. [29], showed that new superferromagnetic tracers improve the resolution and sensitivity of MPI by more than 10-fold. These particles show ferromagnetic behaviors such as coercivity and remanence, but the switch from positive to negative saturation is much steeper than superparamagnetic nanoparticles. Fig. 2 demonstrates this by showing an M-H curve similar to ferromagnetic materials, showing hysteresis that is not seen in superparamagnetic materials. It also shows that the nanoparticles with these properties form chains under applied fields. Tay, et al. [29] found that those chained particles meet the definition of superferromagnetism. They found that magnetization of superferromagnetism could be modeled as a Langevin saturator, a transcendental equation where the output magnetism builds upon the induced magnetism of neighboring SPIOs.

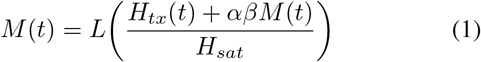

where *H*_*tx*_ is the applied magnetic field, *H*_*sat*_ is 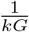, *M* is the magnetization of the chaining nanoparticles, and *β* is a dimensionless parameter that describes the saturation magnetization of the chain. *α* is another dimensionless parameter that describes how closely the nanoparticles in a chain can be packed. Any applied magnetic field to change the chain’s magnetization direction must overcome the magnetic field produced by each nanoparticle’s neighbor. This models the hysteresis, coercivity, and remanence as seen in the superferromagnetic samples.

**Fig. 2.**
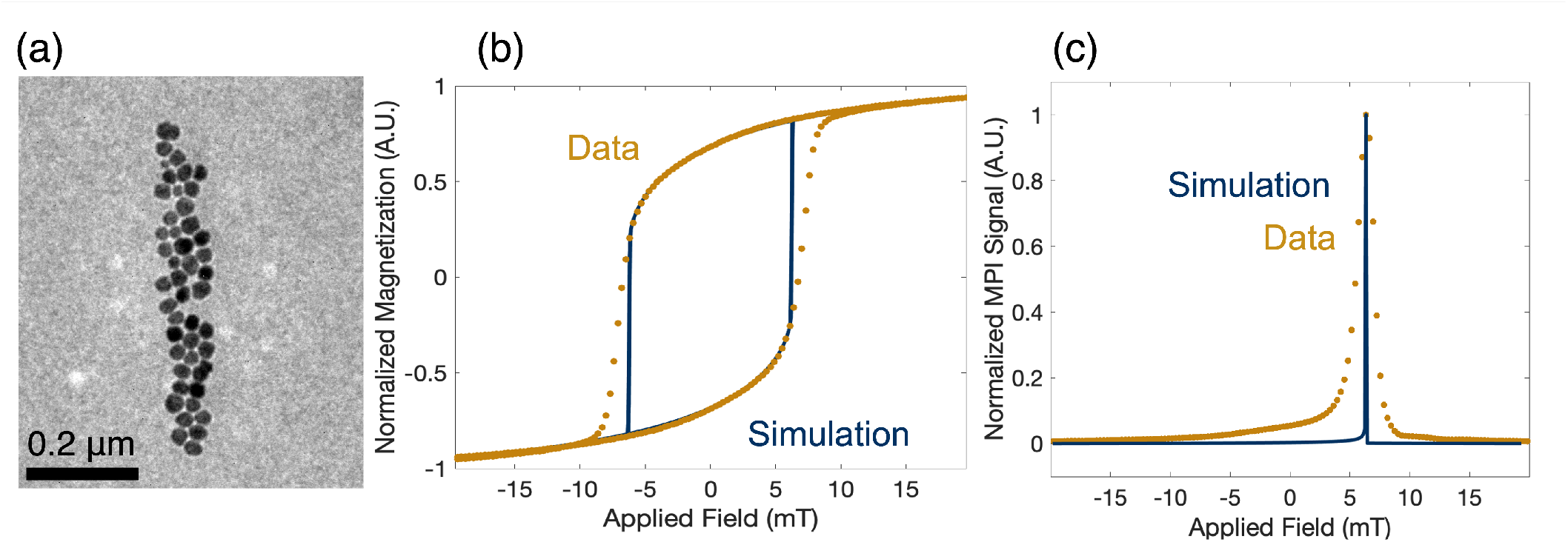
Chaining Hypothesis Experimental Data and Theoretical Predictions (a) TEM Images of Chain Formation in SFMIO samples. SFMIO samples were exposed to 16 mT magnetic fields. Langevin physics predicts that SPIOs align in the direction of the applied field. Samples distinctly show chain formation in the direction of the applied field. (b) Experimental data and Theoretical Predictions of SFMIO M-H curves. The SFMIO curve shows both coercivity and remanence characteristic of ferromagnetic materials. Its behavior is correctly modeled by a Langevin “saturator” element as detailed in [29]. (c) Theoretical Predictions and Experimental data of SFMIO PSFs. There is strong agreement between the theoretical predictions and the experimental data.

This function also alludes to the process of chain formation because *M* (*t*) increases in magnitude as the particles draw closer to one another. Assuming that the conditions of potential aggregation of nanoparticles are met, the equation begins with a nanoparticle feeling the magnetic field of its neighboring nanoparticle. As the neighboring nanoparticle drifts closer, *β* increases, leading the nanoparticle’s magnetization to increase, increasing the draw of the neighboring nanoparticle, and so on.

#### 1D Chain Formation

The hypothesis of chaining nanoparticles naturally brings new predictions as a consequence. Almost all previous works of MPI assumes that SPIOs are non-interacting. Tay et al. [29] posit that superferromagnetic behavior is caused by nanoparticle assemblies, chains of SPIOs.

If this is true, then I hypothesize these four corollaries.

1. If nanoparticles are below a certain concentration, no superferromagnetic behavior should be observed.
2. The formation of chains should depend on the viscosity of the solvent that the particles are in, even for Ne’elian nanoparticles.
3. Superferromagnetic behavior should be observed after time exposed to magnetic field, called the polarization time.
4. If particles are not polarized past some threshold, presumably the coercive threshold, then superferromagnetic behavior should not be observed.

If these predictions are true, the case for chain formation of SPIOs being responsible for superferromagnetic behavior is much stronger.

It is essential to understand the conditions that lead to the observed experimental results of chain formation. To this end, we developed a 1D model that finds how chain formation time scales with viscous drag, particle size, and initial concentration. This is a simple model that has been derived before, but we found it particularly useful for understanding the chain formation process [34].

In this model, each superparamagnetic nanoparticle has aligned with a constant applied magnetic field. If each particle is close enough to feel the other’s induced dipole moment, then a magnetomotive force is formed, drawing them closer together. The Stoke’s drag of the viscous surrounding fluid resists the motion. With typical SPIOS, the Stoke’s drag on each particle is enough to dominate its induced magnetomotive force.

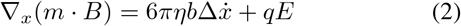

This equation solves for the time it takes for the distance between the two nanoparticles to go to zero, where *m* is the magnetic moment of the particles, *q* is the electric charge of the particles, *η* is solvent viscosity, and *b* is the hydrodynamic radius of the two particles. In the case of our experimental results, the nanoparticles are in a non-polar solvent and thus have no surface charge. Therefore, the electric field term can be disregarded, leading to a simple solution for the distance between the two nanoparticles as a function of time

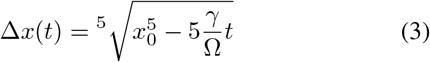

where *x*_0_ is the starting distance the particles are apart from one another and the analogue to concentration, *γ* is 3*µ*_0_*m*^2^*/π*, where *m* is the magnetic moment of the particle. Ω is 12*πηb* where *η* is solvent viscosity and *b* is the particle’s hydrodynamic radius. The solution to equation 3 indicates that the chain formation time, which is the time it takes for particles to attract one another, depends most strongly on the concentration of the particles in solution. There should also be a dependence on viscosity: the more viscous the solution, the longer it takes for chains to form.

## III. Materials and Methods

### Nanoparticle Synthesis and Properties

#### Phantom Scans

Phantoms were made with 300-micron diameter holes spaced 2.35 mm apart. They were filled with SFMIO or Vivotrax tracers. MPI scans took place on a custom-built vertical bore 2D Projection MPI Scanner with a 5.9 T/m gradient field. Both SPIOs and SFMIOs underwent the exact same pulse sequence. Nanoparticles were excited at a transmit frequency of 20.225 kHz and a transmit amplitude of 20 mT. The scanning phantom was mechanically translated in the z-direction at 0.5mm increments. The FOV was 5.1 cm by 4.6 cm, with a scan time of 5 min. Images were reconstructed by averaging the positive and negative scans. They were then filtered with 1D gaussian filter to dehaze the images.

#### SFMIO Synthesis

Prior work detailed and experimentally confirmed the process of creating single-core magnetite nanoparticles [35]. This adapted method had two steps: first, the creation of an iron-oleate precursor, and second, the thermal decomposition of the precursor to form magnetic nanoparticles. The synthesis procedure was as follows: first, a stoichiometric amount of iron acetylacetonate and oleic acid (3.3 g and 15 mL, respectively) were placed in a reaction flask with stirring at a rate of 300 rpm. The flask was then placed in a molten solder bath at a temperature of 290^*◦*^ C. The temperature of the bath was then raised to 320^*◦*^ C. 30 min after being placed in the bath, the flask was removed and allowed to come to room temperature. The next day, the prepared iron-oleate precursor was used to synthesize the magnetic nanoparticles. First, the precursor, at 0.62 M, was diluted with 1-octadecene to create a 0.22 M mixture. The diluted mixture was then loaded into a syringe. It was then dripped into a reaction flask containing 5.1 mL of octadecene for 25 minutes. The solder bath was stabilized at 360^*◦*^ C and the sample within the reaction flask was stirred vigorously at a rate of 300 RPM. After cooling overnight, the resulting mixture was then washed in hexane in a 1:1 mixture, then centrifuged at 14 kRCF for 30 min. The resulting supernatant was discarded and only the precipitated particles were saved. This washing process was repeated multiple times.

#### Nanoparticle Characterization

The precise mass of iron in the SFMIO sample was determined using Perls’ Prussian blue reaction as described in [36]. In this process, 40 *µ*l of SFMIOs were first digested in 1 mL of 12M HCl. After digestion, the resulting solution was mixed with 100 *µ*l of 5% postassium ferrocyanide. Absorbance was measured in a UV-Vis Spectrophotometer at two wavelengths, 680 nm and 730 nm. The absorbance of the SFMIOs was then compared to a calibration curve made through the same process that used SPIOs of a known concentration of ferucarbotran (Vivotrax). TEM images of chained SPIOs were made by placing 3 *µ*l samples on formvar-coated copper grids that then dried under static 22 mT applied fields for 5 h. The chains were then confirmed by transmission electron microscopy (JEOL 1200EX, 80 kV). For the viscosity experiments, the solvent viscosity was determined by referring to [37]. In this paper, proportional mixtures of hexane and squalane were created and their material properties measured. We first dried SFMIO samples in hexane. We then resuspended said samples in mixtures of hexane and squalane. We then performed a nonlinear least squares fit on [37] data and back calculated the resulting viscosity of our samples. This gave us an estimate of the solvent viscosity.

#### Resolution and Sensitivity Calculation

Spatial resolution is calculated, using the Houston Criterion, from the Full Width Half Maximum (FWHM) of the image of a point source, which is often called the Point Source Function or PSF [38]. Resolution in MPI is converted from a magnetic resolution (mT) to a spatial resolution (mm) by dividing the result by the 3D scanner’s gradient field strength (T/m). The Berkeley 3-D MPI Scanner has a gradient of 7 T/m, leading to 1 mT becoming a spatial resolution of 140 *µ*m. MPI Signal strength (mV/mg Fe) was calculated by dividing the voltage of the received signal by the iron mass of the tracer. That number is then normalized by the concentration-normalized MPI signal measurement of Vivotrax.

### Pulse Sequence to measure Chain Formation Time

The SFMIOs were tested on our arbitrary waveform relaxometer (AWR), as described by prior work from the lab [39]. The AWR’s non-resonant coil design enables the creation of custom pulse sequences, hence, arbitrary. The AWR can reconstruct the magnetization response such that a 1D point-spread function (PSF) characterizes the entire sample within the device. The chain formation was measured using a unique pulse sequence designed to quantify the change in PSF as a function of the length of time a pre-polarizing pulse was applied to the sample. The pulse sequence is repeated 1 ms long pre-polarizing pulses (TP) followed by 20 kHz (TR) readouts. This sequence (pre-polarizing pulse + readout) was repeated continuously for a total sequence length of 1.4 s.

## IV. Discussion and Results

IV-A demonstrates the theoretical predictions of the Langevin saturator model versus experimental data. IV-B shows the change in SFMIO PSF as a function of time exposed to an applied field with a comparison of Vivotrax’s response. IV-C shows how the SPIO to SFMIO transition is affected by the amplitude of the applied field. IV-D demonstrates that the measured PSF can be linearly decomposed as a transitions between a population of SPIOs to a population of chained SPIOs, aka SFMIOs. IV-E shows that the transition from SPIO to SFMIO is affected by both the concentration of the nanoparticles within the sample and the viscosity of the solvent the nanoparticles are in.

### A. Langevin Saturator Theoretical Prediction of Experimental Data

The Langevin saturator model in Eq. 1 predicts step-like transitions from positive to negative saturation or vice versa. It also predicts coercivity and remanence, much like a ferromagnet. Fig. 2 shows experimental data and theoretical modeling of the SFMIO M-H curve. These predictions were experimentally verified, showing that SFMIO behavior is accurately modeled by the Langevin saturator model. Deviations from the predicted behavior during the process of transitioning from one saturation to another are thought to be due to coercivity dispersion. Coercivity dispersion is when a sample has multiple coercivities, due to different lengths of chains.

### B. Transition from SPIO to SFMIO Behavior as a Function of Pre-Polarization Time

Our results in Figs. 5 and 6 clearly show the difference in SFMIO PSF properties as a function of pre-polarization time. In Fig. 5, the change is shown in 3 PSFs at different pre-polarizing times. The most dramatic change in resolution occurs within 25 ms of exposure to a pre-polarizing field. This fast change in transition is supported by Fig. 6, where the transition in both resolution and peak signal of an SFMIO sample occurs within the first 50 ms. By 1.4 s of pre-polarizing time, the final PSF has a resolution of 1 mT and a 6-7x increase in peak signal. This leads to a total improvement in resolution by 10x and an 6-7x increase in signal strength. In contrast, Vivotrax, undergoing the same pulse sequence, stayed constant in both resolution and peak signal.

**Fig. 3.**
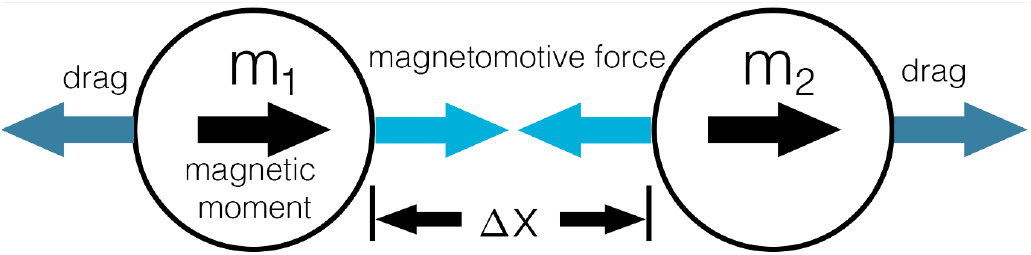
1 dimensional free body diagram of two nanoparticles affected by one another’s induced magnetic field. As their magnetic moments align with the spatially constant applied magnetic field, an attractive magnetic field is induced in each nanoparticle. This induced magnetic field does vary spatially, and therefore, it produces a magnetomotive force. 1D Stoke’s drag resists the magnetomotive force that draws the two nanoparticles together.

**Fig. 4.**
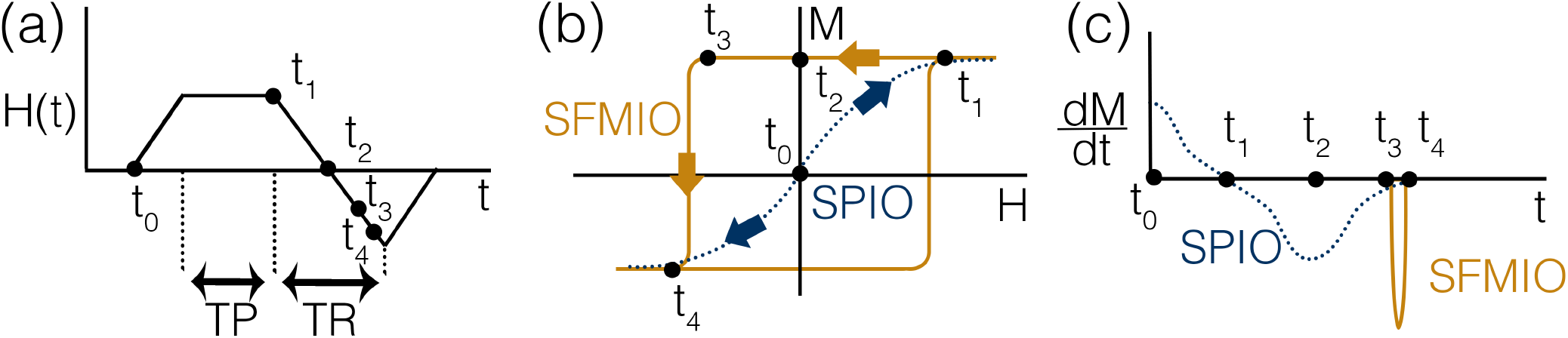
This pulse sequence allows us to uniquely observe the chain formation time with excellent SNR. (a) Chain Formation Pulse Sequence. From ***t***_**0**_ to ***t***_**1**_, the AWR is ramped up to a constant applied field and a pre-polarizing pulse is applied, the length of which is TP. ***t***_**1**_ to ***t***_**4**_ shows the readout time, TR, which describes the length of time the applied field goes from maximum to minimum amplitude. (b) M-H curve showing SPIO and SFMIO magnetization as a function of applied field. If the sample shows SPIO behavior, the SPIO magnetization curve is followed. If the sample shows SFMIO behavior, the SFMIO magnetization curve is followed. c) 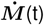 plot showing the inverse of what the receive coil measures. From ***t***_**0**_ to ***t***_**1**_, an SPIO PSF is seen. No SFMIOs have been formed yet. If no chains are formed after TP, SPIO behavior will be seen during TR, ***t***_**1**_ to ***t***_**4**_. If chains do form during TP, TR will show SFMIO behavior, where the steep change in magnetization change from ***t***_**3**_ to ***t***_**4**_ in (b) is responsible for a sharp resolution and increase in SNR seen in (c).

**Fig. 5.**
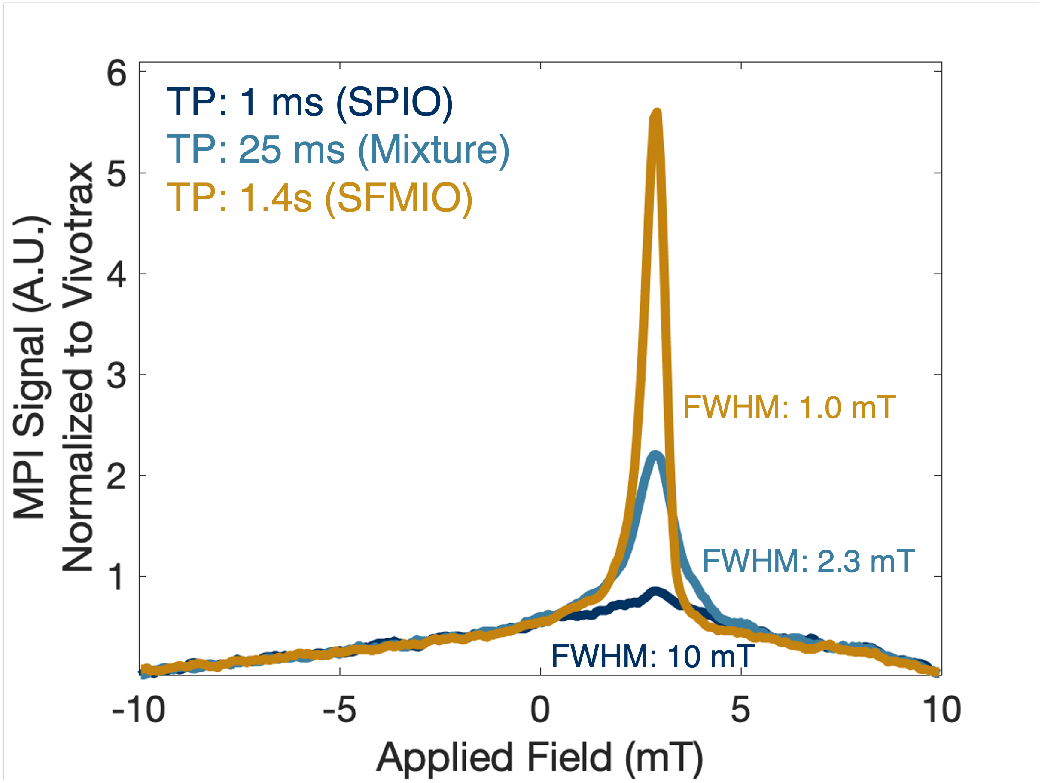
PSF of SFMIO sample as a function of pre-polarization time. As the amount of time the sample is exposed to an applied magnetic field increases, the MPI signal and magnetic resolution improves accordingly. The PSF at a pre-polarization time of 1ms demonstrates SPIO behavior. The PSF at a pre-polarization time of 25 ms shows a 5x improvement in resolution and a 2x increase in MPI signal. We consider the PSF at 25ms to show a mixture of SPIO and SFMIO behavior. After 1.4 s have passed the MPI signal has increased by 6-fold and has a 2-fold improvement in resolution. This PSF shows fully SFMIO behavior, with a total 6-7x increase in MPI signal and a 10x improvement in magnetic resolution.

**Fig. 6.**
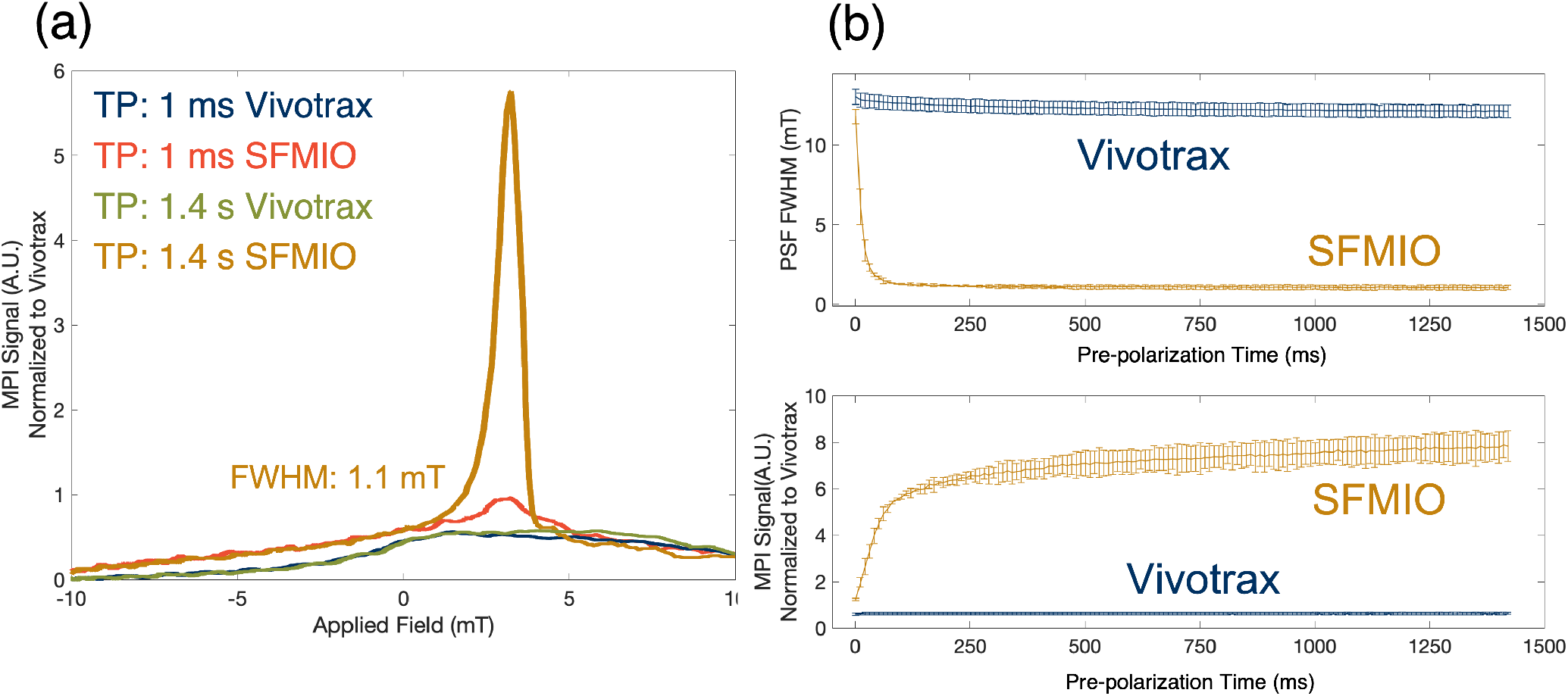
Change in MPI PSF and MPI signal as a function of pre-polarization time (TP). (a) The SFMIO PSF from a 1ms TP to a 1.4 s TP shows a complete change in magnetic resolution as compared to the Vivotrax PSF. (b) The SFMIO sample undergoes a 10x improvement (11.8 mT to 1.1 mT) in magnetic resolution as a function of TP. The Vivotrax sample stays constant in the same pulse sequence with a magnetic resolution of 12.5 mT. The SFMIO MPI signal undergoes a 7x increase as a function of TP while the Vivotrax MPI signal stays constant.

This change in particle resolution and peak signal as a function of time exposed to an applied field affects MPI pulse sequences by indicating that, for a very short amount of time, each sample should be polarized into chain formation before implementation of a pulse sequence. Because the time that the polarization takes for most samples is within the millisecond range, this should not be a difficult requirement to meet.

### C. Transmit Amplitude Threshold of SFMIO Signal

Fig. 7 provides additional support to the Langevin saturator model by confirming the role of transmit field amplitude in the chain formation process. When the transmit amplitude is below this threshold (2 mT), no SFMIO behavior is seen. Fig. 4 shows that, unless the some transmit amplitude threshold is reached, only an SPIO response is seen. However, when the transmit amplitude was greater than that threshold (12 mT), SFMIO behavior was seen.

**Fig. 7.**
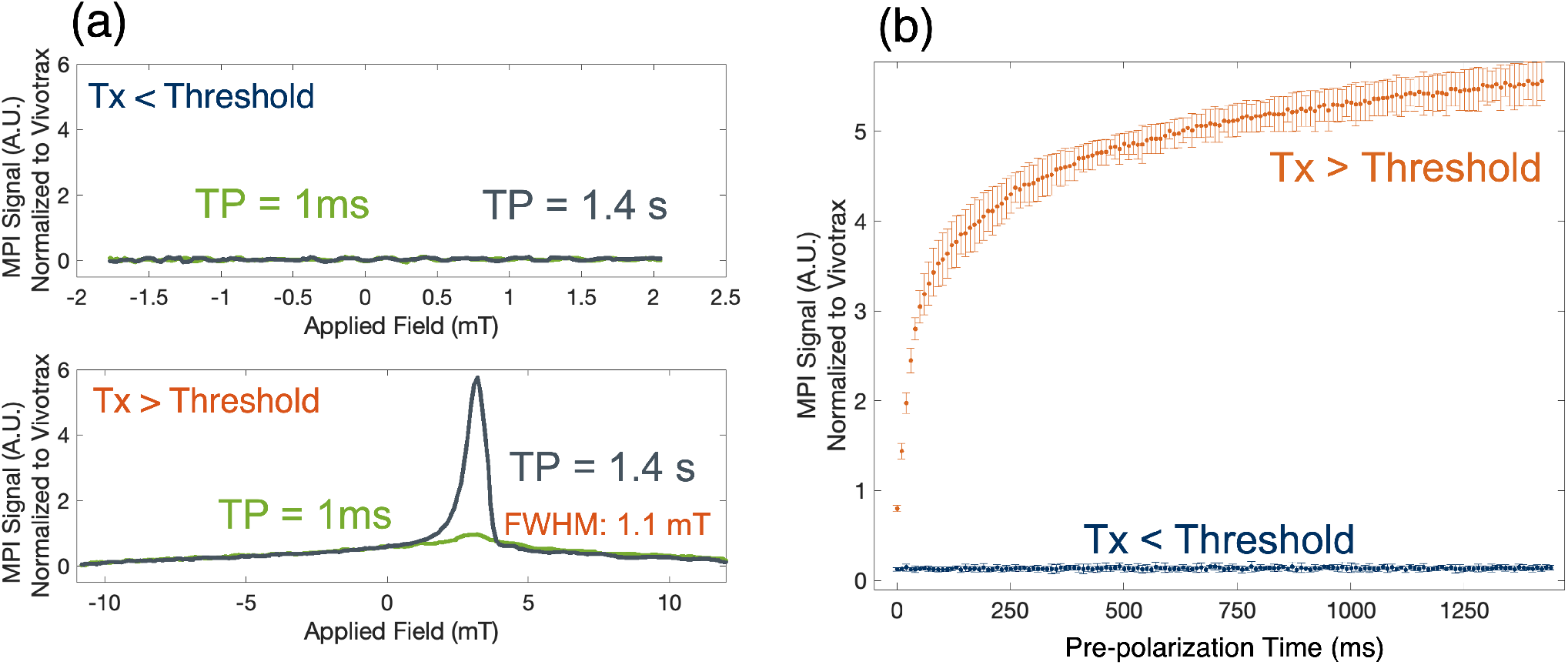
In this experiment, the pulse sequence shown in Fig. 4 is repeated. However, the transmit amplitude of the applied field is varied in two different cases, one above the applied field threshold as shown in Fig. 4 (b), and one below the applied field threshold. The chaining hypothesis indicates that SFMIOs will only “flip” and show SFMIO behavior if the transmitted magnetic field is above the particle’s coercivity threshold. This plot shows that to be a possibility, where a transmitted field of 2 mT shows an MPI signal of less than 1, while a transmitted field of 12 mT shows the expected SFMIO behavior, a reduction of 10x resolution and an increase in MPI signal.

The existence of a threshold for SFMIO behavior means that transmit amplitude must be greater than either a sample’s coercivity or saturation field (*H*_*sat*_). We would need nanoparticles where *H*_*sat*_ is smaller than the coercive threshold of the chain to test the mechanism for the threshold thoroughly. We currently have only synthesized particles that have a coercivity threshold less than *H*_*sat*_. The threshold observed is most likely a coercivity threshold.

We found experimentally that each “batch” of SFMIOs synthesized has different coercive behaviors. While this is the case, it is possible that each SFMIO batch needs to be characterized before its use in MPI pulse sequences. However, SFMIO synthesis is still very much in early stages. There is room for improvement in the synthesis process.

### D. Linear Decomposition of Signal

Analysis of the PSF change as a function of pre-polarization time as seen in Fig. 5 shows that transition from SPIO to fully chained SFMIOs does not occur immediately and goes through transition phases. Intuitively, this seems very reasonable. Each TP and TR are repeated, with a total scan length of 1.4 s. From the first TR to the last TR, the amount of iron nanoparticles within the sample stays constant. The SPIOs chain as they are exposed to pre-polarizing pulses. This chaining occurs over some period of time. Therefore, it is reasonable to say that there are two populations of particles within the sample, chained and unchained.

In this analysis, we make three assumptions. The first is that the very first PSF, barely exposed to the pre-polarizing pulse, shows completely SPIO behavior and has no chained SPIOs. The second assumption is that the final PSF that has been exposed to 1.4 s of pre-polarizing time shows completely SFMIO behavior and has negligible free-floating SPIOs giving signal. The third assumption we make is that, in the sample, the nanoparticles can either be free-floating or chained. We then showed in Fig. 8 that we can linearly decompose the results of our pulse sequence into two populations - free or chained particles. Note that the point where the SFMIO signal begins to dominate aligns with Fig. 6’s change in resolution and peak signal.

**Fig. 8.**
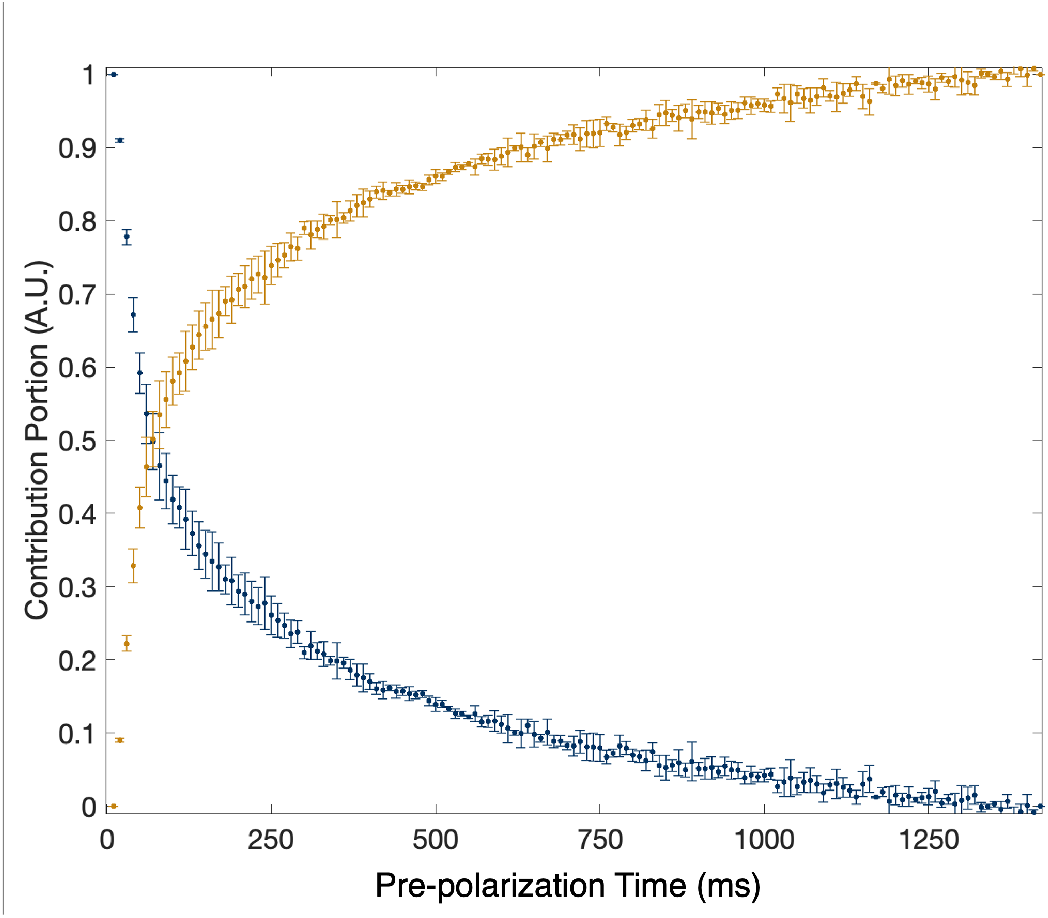
Linear decomposition of sample transition from fully SPIO to fully SFMIO behavior. This shows the transition from 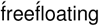 SPIOs to chained SPIOs, aka SFMIOs. Note that the transition from SPIO to SFMIO domination occurs at the same time as Fig. 6ś noticeable change in Peak Signal and resolution. This is from the same data set.

### Concentration and Solvent Viscosity Dependence of Chain Formation Time

Eq. 3 indicates that the time it takes for SPIOs to form chains will depend on the concentration of iron in the sample and the viscosity of the solvent the nanoparticles are in. The initial interparticle distance, *x*_0_, relates to the initial concentration by 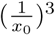. *x*_0_ then has a fifth root relationship with the chain formation time. Fig. 9 shows how even a modest change in concentration (as seen in the bottom plot) dramatically boosts the magnetic resolution if the concentration permits chain formation. For example, a 3x reduction in concentration (from 2.7 to 0.7 mV/mg Fe) leads to virtually no change in the resolution, but another drop of 0.3 mg/ml, almost completely turns “off” any SFMIO behavior. Tay, et al. [29], hypothesized that biocompatible encapsulation, much like micelles, should be used to enforce a high local concentration of SFMIOs and a low systemic concentration for biosafety. These concentration results show that threshold concentrations must be exceeded such that SFMIO behavior is still seen within biocompatible encapsulation.

**Fig. 9.**
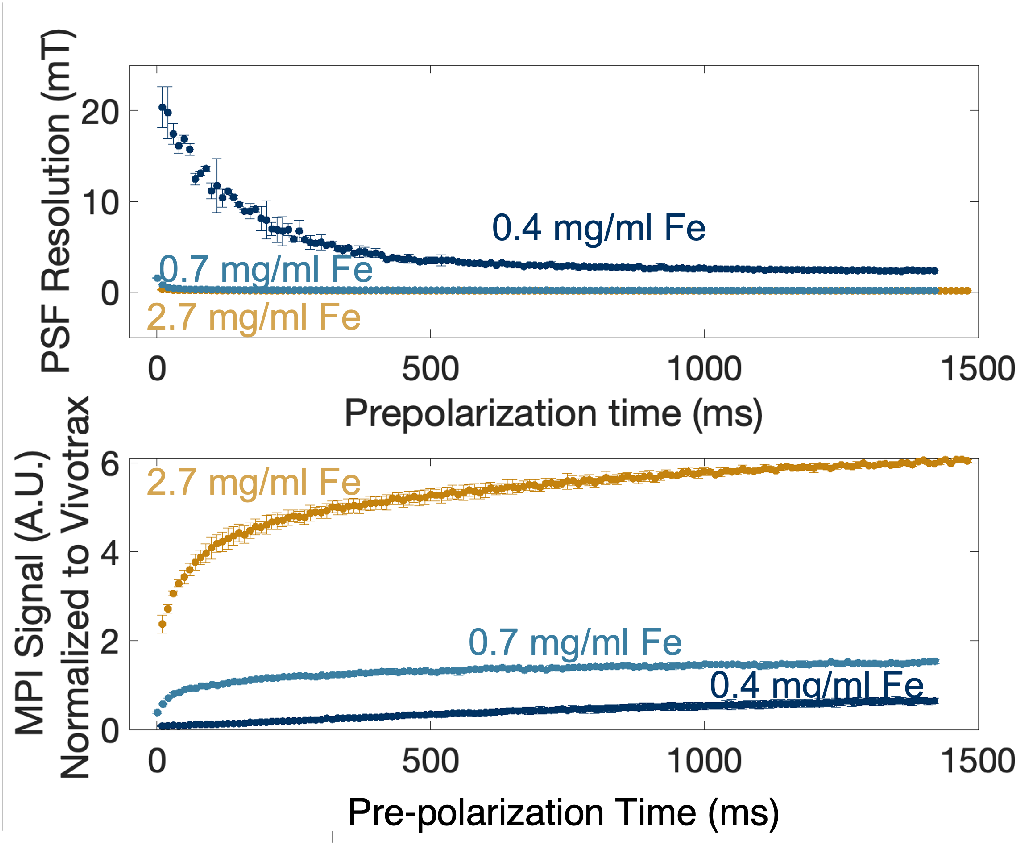
Change in SPIO to SFMIO transition as a function of particle concentration. In the top plot, a small change in behavior occurs between 2.7 mg/ml Fe and 0.7 mg/ml Fe. However, a very large change in SFMIO behavior occurs with a 0.3 mg/ml difference in sample concentration, with the 0.4 mg/ml sample showing a much less steep transition between the first magnetic resolution measurement and the last. The bottom plot demonstrates that the peak signal of each sample is affected by the concentration of the particles, with the lowest concentration showing very little change in peak signal as a function of pre-polarization time at all. Note that while the 0.7 mg/ml sample shows very little change in magnetic resolution from the 2.7 mg/ml sample, its peak signal is 5x lower in SNR. This is, presumably, from the reduction of iron in the sample.

Fig. 10 shows how the change in solvent viscosity affects SFMIO behavior. Solvent viscosity has a crucial effect on the SFMIO behavior; if the fluid viscosity is too high, then Stoke’s drag will dominate the magnetomotive force and hence chain formation will occur very slowly. In the 1D Chain formation model, the solvent viscosity should have a fifth root relationship with the chain formation time. We saw that as solvent viscosity increased, the chain formation time increased. Note that the final resolution of each sample was also worse. This underlines how important it will be to ensure low viscosity fluid inside biocompatible encapsulations.

**Fig. 10.**
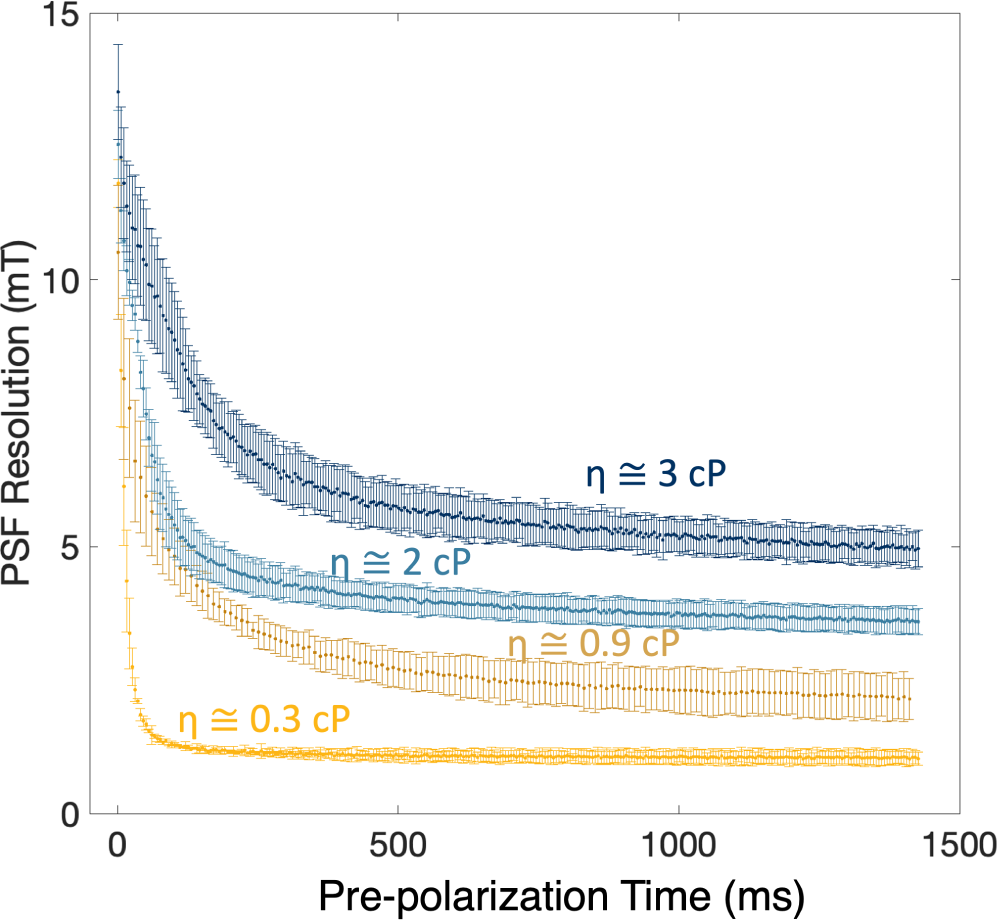
Plot of PSF magnetic resolution vs. pre-polarization time for various viscosities. The change in resolution from SPIO to SFMIO behavior seems to occur around the same time, which is not predicted by the chain formation model. Note that the final magnetic resolution of each sample lowers as the solvent viscosity increases. There does not seem to be any further decrease in magnetic resolution as a function of pre-polarization time.

## V. Conclusion

In conclusion, we have demonstrated that SFMIO behavior does not occur immediately and that a transition between SPIO and SFMIO behavior occurs as a function of exposure to magnetic fields. This result supports previous work that showed that chains of superparamagnetic nanoparticles were responsible for order-of-magnitude improvement in MPI sensitivity and resolution.

MPI is a promising pre-clinical *in vivo* imaging modality. It has many strengths, e.g., no signal depth dependence, linear positive contrast, non-radioactivity. However, to this point, its clinical implementation has been held back by its poor resolution in small animal scanners. Superferromagnetic iron oxide nanoparticles are a new tracer for use in MPI that alleviate this resolution issue by 10x, along with an increase in sensitivity.

These results in this chapter will inform future work on biocompatible encapsulations for SFMIOs and pulse sequence design. Pulse sequences must have transmit amplitudes that exceed the transmit amplitude threshold of the sample SFMIOs. Pulse sequences must also include short periods of time for chains to fully polarize. The biocompatible encapsulation for the particles themselves must have concentrations within that are high enough to see SFMIO behavior, and the solvents inside the biocompatible encapsulation must be as low viscosity as possible. These conditions will help enable SFMIOs to be successfully used in MPI scans.

## Acknowledgment

Steven M Conolly is a co-founder of an MPI company, Magnetic Insight, and holds stock in this company. The authors declare no other conflict of interest.

The authors thank the staff of the Electron Microscopy Lab and Biomolecular Nanotechnology Center – QB3 at UC Berkeley for the usage of their equipment.

